# Machine Learning Prediction of HIV1 Drug Resistance against Integrase Strand Transfer Inhibitors

**DOI:** 10.1101/2025.04.25.650610

**Authors:** Ching Yu Chui, A.W. Edith Chan

**Affiliations:** Division of Medicine, Cruciform Building, University College London, Gower Street, London WC1E 6BT, UK

## Abstract

Infection caused by the human immunodeficiency virus (HIV) can be effectively treated using antiretroviral therapy (ART). One such therapy involves drugs that target the HIV integrase. However, this has resulted in the development of resistant associated mutations (RAMs). This investigation aims to create a machine learning model to classify an input protein sequence of HIV integrase as ‘resistant’ or ‘non-resistant’ towards the five approved integrase strand inhibitors (INSTIs). The training data consists of protein sequences along with the associated biological features of each residue: its presence in the drug binding site, secondary structure, solvent accessibility and mutation frequency. A logistic regression model was developed and from this model, key residues which contribute towards drug resistance were identified, including several known RAMs. The model performance was on a par with other similar studies that used for classifiers with more complex architectures. The approach described here could be adapted to other resistance-prone diseases.

## INTRODUCTION

Recent advances in applying machine learning to biological problems can offer new perspectives for HIV drug resistance. While it is possible to simply express the protein using the viral sequence within a patient and measure drug resistance using a phenotypic assay, this can be expensive and time consuming. Therefore, a genotypic approach is preferred.

Predicting drug resistance from sequence data can be viewed either as a problem of regression or classification. For the regression approach, the goal is to train a machine learning model to return an indicator of drug resistance, for instance IC_50_ fold change. Different types of machine learning models have been used successfully in analysing aspects of HIV resistance data. For instance, an artificial neural network, a type of regression model, was used to predict HIV resistance against a range of protease and reverse transcriptase inhibitors.^1^ Similarly, a support vector machine model was created to calculate the probability of drug resistance.^2^ Two other models, random forest and k-nearest neighbour, have also been implemented to predict resistance.^3^ By using regression models, numeric data is returned.

This allows the user to examine the extent of resistance, which can better inform drug administration.

Alternatively, the problem can be viewed as one of classification. The goal of the machine learning model is to classify a given sequence as ‘resistant’ or ‘non-resistant’ to specific drugs. This requires a certain threshold value, for instance in the form of phenotypic data, to be determined for classification. In one example, logistic regression has been used to analyse HIV reverse transcriptase sequences, and revealed the importance of using cross-resistance information in determining resistance against nucleotide analogues.^4^

Recently, several types of deep learning classifiers have been used to analyse HIV resistance data across 18 different ART drugs.^5^ Deep learning models are a subset of machine learning models using multiple layers of neural networks to achieve higher learning capabilities. However, the ‘black box’ nature of these models can reduce their interpretability. To overcome this shortcoming, one study used tools that improve model interpretability and successfully highlighted relevant features.^5^ This study not only showed the potential of deep learning models in performing sequence data analysis, but also in the identification of new RAMs.

In another study, three binary classifier models were used to identify new RAMs: multinomial naïve Bayes, logistic regression and random forest. The classifier models in this study performed well in the presence of RAM-containing sequences and poorly in their absence. Six potential new minor RAMs were identified and found only increase resistance when present with known major RAMs.^6^

All these studies have yielded encouraging results regarding the HIV drug resistance landscape, supporting the concept that machine learning models are powerful tools for studying sequence data. However, a possible limitation is the lack of a biological perspective. These models were trained on phenotypic data and features only containing amino acid positions and mutations, have not reflecting the reality that physiologically, proteins are three-dimensional macromolecules in a complex solvent environment. While much useful information can be extracted from the protein’s sequence, it is its three-dimensional fold that determines its biological function.

In this study, we are interested in drug resistance to integrase strand inhibitors (INSTIs). Figure 1 shows a list of currently approved INSTIs. For the machine learning model worth including is feature the position of a given residue relative to the drug binding site, as INSTIs are competitive inhibitors directly binding near the protein-DNA interface. Therefore, a mutation within or near the binding site is more likely to influence drug binding and therefore resistance. After model training, it would be interesting to see if important features map to the drug binding site. Using whether the residue occurs in the binding site can detect the significance of any other binding sites that may contribute to drug resistance towards INSTIs.

**Figure 1:**
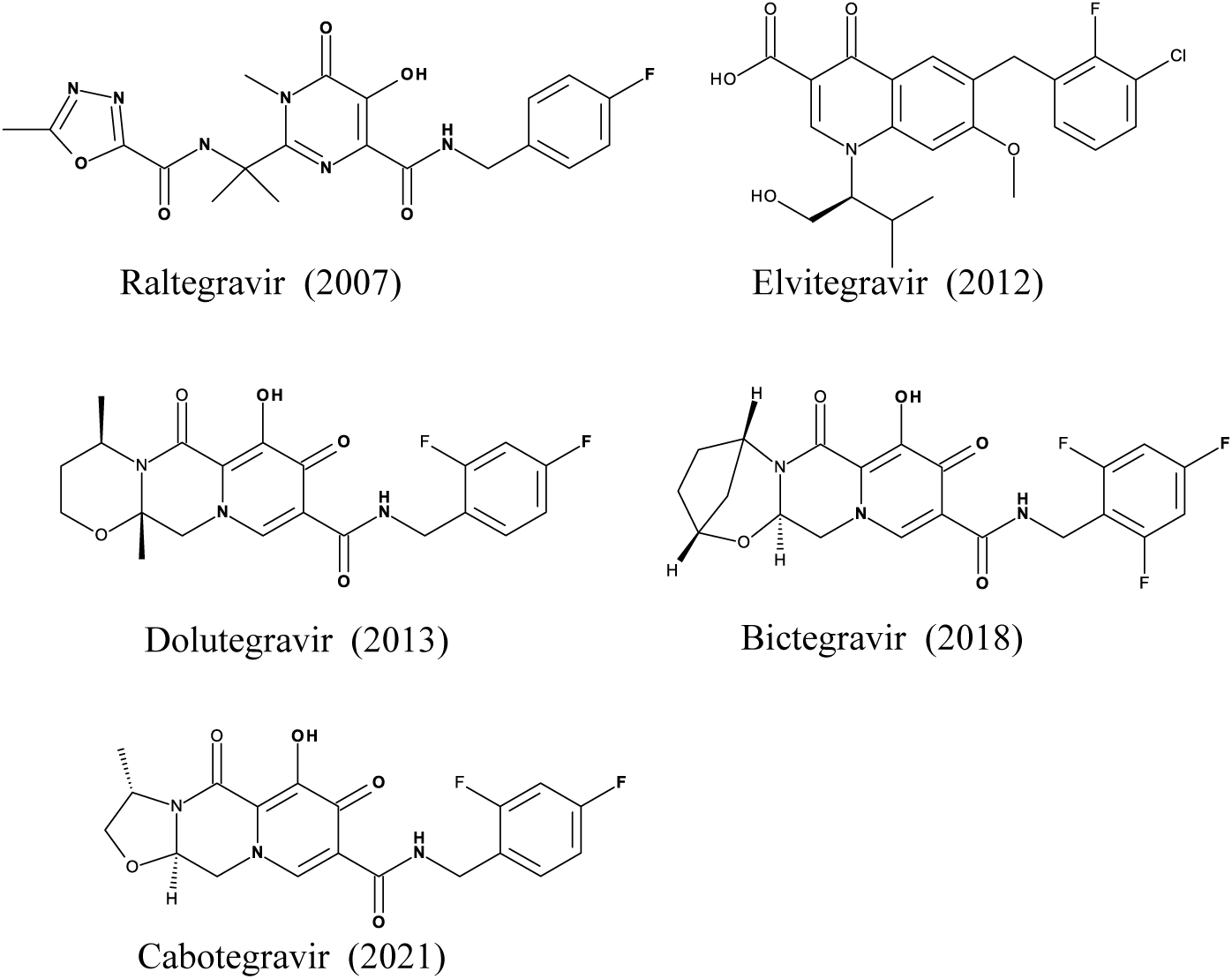
Five currently approved HIV integrase inhibitors along with their 2D-chemical structures and year of FDA approval in parenthesis.

Another key consideration is the solvent accessibility of residue. This is important as it determines if it is openly accessible indicating it to be on the protein surface or near a binding site; inaccessible residues tend to be buried. A recent computational study^7^ performed molecular dynamics and free energy perturbation on a set of inhibitors bound in the protein binding site. In their findings, residue Q148 of integrase was identified as important in affecting the distribution of water molecules in the protein’s central pocket causing Q148H/K/R to be major RAMs.^7^ Given this information, it may be useful to measure solvent exposure at each residue. This will function as an indirect measure of the chance of being affected by free energy change caused by water for residues near the binding site.

Additionally, the mutation frequencies of each residue should be considered. Identifying which residues have a higher tendency of mutation in both treatment-naïve and treatment-experienced populations can provide an insight into which RAMs are more likely to arise and also identify RAMs that are induced by treatment. For instance, natural polymorphism is common at S119 and has no effect on its own. However, S119R can increase drug resistance conferred by known major RAMs such as Q148H.^8^

INSTIs are a class of drugs not frequently discussed in HIV drug resistance studies involving machine learning. Given that most HIV RAMs are already well-established and studied, the aim here is to create a binary classifier that determines whether a given sequence will be resistant towards the five currently approved INSTIs. While the main features in our study remain the individual amino acid positions, the biological parameters described above will also be incorporated into our model, namely whether the residue is in the drug binding site of integrase, its local secondary structure, solvent accessibility and mutation frequency.

## MATERIALS AND METHODS

### Dataset

Integrase sequence data, labelled with resistance data, was obtained from the Stanford University HIV drug resistance database.^9^ All sequences containing resistance data for raltegravir, elvitegravir, dolutegravir, bictegravir and cabotegravir were selected. This resulted in 5143 sequences. The data were filtered such that only data obtained from the PhenoSense assay were used so the results could be directly comparable. Resistance data were given in the form of resistance fold change, which can be calculated by:

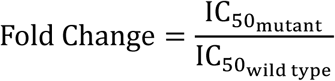

The value of the threshold fold change for resistance was 3.5 as defined in the database.^9^ This gives 2720 sequences.

Next, these sequences were aligned with the wild type integrase sequence (NCBI accession: 8W34_A) and trimmed. During trimming, the longest common sequence was used as reference. At the end, all sequences were 288 residues long.

To address any duplicate sequences, the most common duplicate was identified and for use as a negative control in the mutation frequency analysis. In effect, this would diminish the effect of naturally occurring polymorphisms and bring focus to mutations induced by drug administration.

### 3D binding site features

To determine whether residues belong to the integrase binding site, three 3D structures of integrase bound to the largest INSTIs were visually inspected using PyMOL^33^ (PDB: 8fn7^31^, 6puw^30^, 6puy^30^) to ensure full coverage of the binding site. The binding site for each PDB is defined by all atoms within a distance of 6Å from any atom in the bound ligand. All residues participating in the binding site were recorded. The resultant binding site thus would likely encapsulate all key residues involved in INSTI binding.

Secondary structures were predicted using the DSSP program^10^ and verified using the X-ray structure (PDB: 8fn7^31^). Solvent accessibility was calculated using Naccess,^11^ with the default Z-slice thickness of 0.05 Å as a compromise between accuracy and computational speed. The probe size in Naccess was set to the default value of 1.4 Å.

### Mutation frequency analysis

For the mutation frequency analysis, a larger dataset of sequences was used to increase the accuracy of the analysis since no phenotypic drug resistance data was required for this. After extracting the integrase sequences from the Los Almos HIV sequence database^12^, the mutation frequency of each residue was calculated as follows:

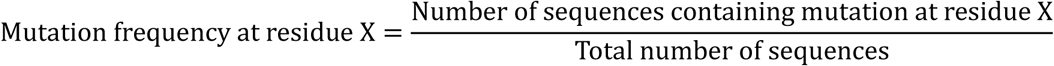

The positions of major RAMs were mapped onto the data to see if there are any trends, such as whether major RAMs are more likely to mutate than other residues. Furthermore, for positions with major RAMs the exact amino acid mutated and the percentages were recorded.

### Encoding the Sequence Data

Sequence data extracted from the Stanford database were strings of letters. To use these sequences as input for the machine learning model, the sequences had to be converted into numeric data, for which one-hot encoding was used, in which each amino acid was transformed into a vector with each row denoting whether the amino acid matches to the one assigned to each row. This approach can address the bias introduced if integer encoding is used instead and ensures higher numbers will not be treated as more important. At each position, further rows were added to the vector to reflect the biological features described above. Binding site residues were assigned in a binary fashion: within (1) or outside (0) the binding site. Secondary structures were integer encoded. Alpha helices were assigned 1, 3_10_ helices 2, beta strands 3, H-bonded turns 4, bends 5 and isolated beta bridges 6. Solvent accessibility and mutation frequency data were directly added into the vectors without further modifications as they were numerical.

### Logistic Regression

Various machine learning models have been used in similar studies of drug resistance. For those using a classification approach, classifier chains^13^ and support vector machines^2^ have been tried. In another case, the performances of multinomial naïve Bayes, logistic regression and random forest methods were compared against one another.^6^ Another approach was to use deep learning architectures. While these might result in higher prediction accuracy, there were several limitations, especially in deep learning architectures, where there is a lack of user-defined constraints, making the data-fitting process more difficult.

In this investigation, logistic regression was chosen as the binary classification model. Due to the limited number of sequence data available labelled with phenotypic resistance data (2720 sequences), deep learning models are unlikely to give accurate results whereas logistic regression has been shown to outperform multinomial naïve Bayes and random forest in predicting drug resistance.^6^ Furthermore, logistic regression has higher interpretability as its output is given in the form of probabilities, which allow the results to be more successfully translated into clinical settings.^6^ In order to address the possibility of overfitting, regularisation was applied. In a model where most features are likely to be unimportant, L1 regularisation is appropriate as it sets small coefficients to zero, thereby lowering model complexity and decreasing prediction error. Therefore, a logistic regression model was trained with L1 regularisation applied.

### Cross Validation

The model was verified using *k*-fold cross validation. In this investigation, there were 2720 sequences in the dataset. Hence, to achieve a compromise between the training data size and the effectiveness of cross validation, the value of *k* was set to be 10.

The two compatible solvers with L1 regularisation, SAGA^14^ and LIBLINEAR,^15^ were chosen and compared against one another. A range of regularisation strength (*λ*) values were tested under both solvers.

The best combination of parameters giving the lowest error were identified before moving on to model testing. The dataset was subjected to an 80/20 split, where 80% of the dataset would act as the training set and the remainder as the test set. From there, model diagnostics could be produced.

## RESULTS

For the machine learning model, only unique sequences were analysed. After removing all duplicate sequences, there were 1103 sequences left from the original 2720 sequences in the dataset. The 80/20 split of the dataset resulted in 882 sequences in the training set and 221 in the test set.

### Binding site residue identification

The binding site for each PDB is defined by the bound ligand, including all atoms to a distance of 6Å. The drug binding surface of each PDB structure was identified as shown (Figure 2).

**Figure 2.**
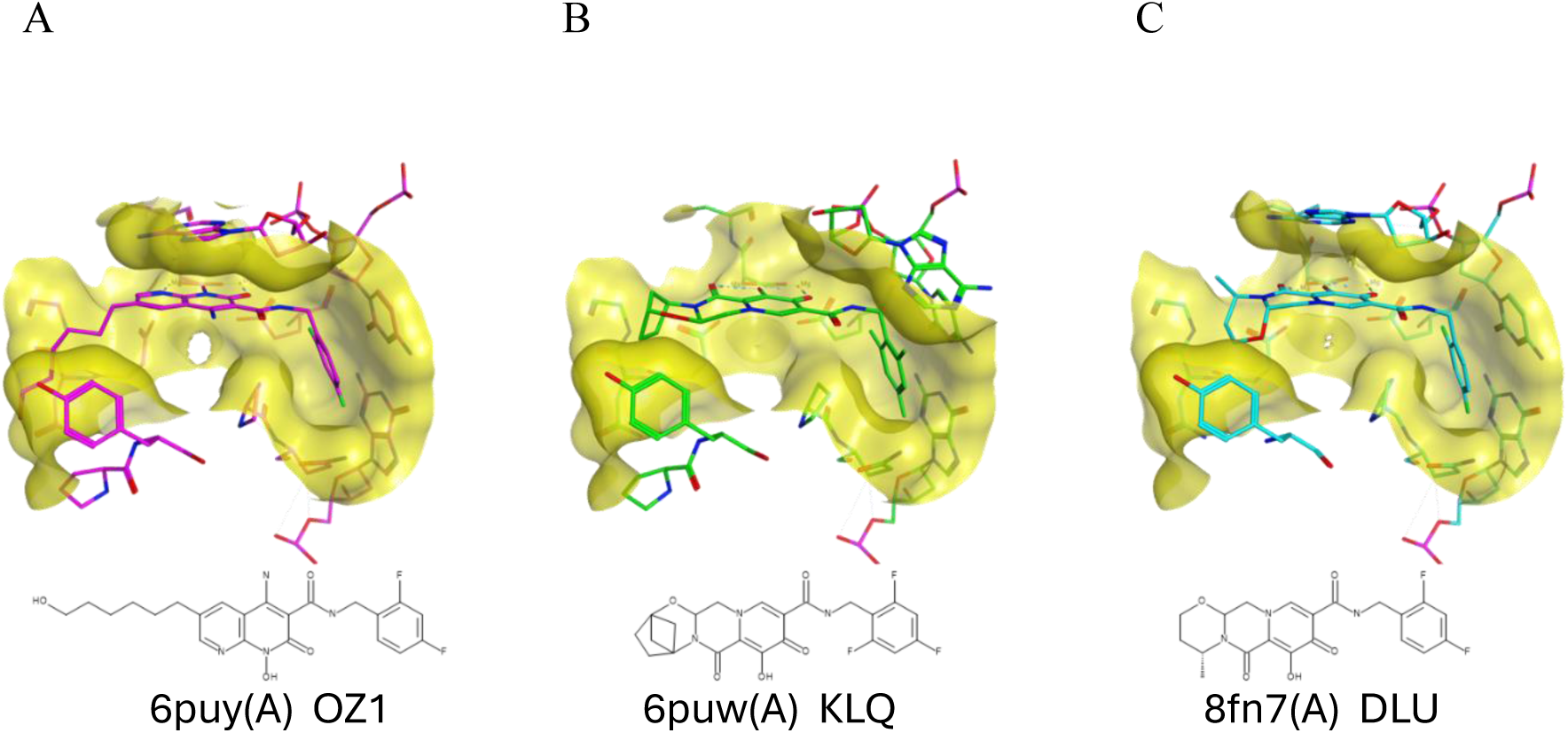
INSTI binding surface drawn with MOE^34^ using its own bound ligand and projected onto its own PDB. A) an experimental INSTI XZ426, B) bictegravir, and C) dolutegravir. The protein chain used is in parenthesis, while the last 3-letter code is the ligand code in the PDB.

The PDB structures were viewed in PyMOL^33^ with residues within its 6Å distance cut-off highlighted. The residues involved in any on of the three of these binding sites were recorded and treated as part of the integrase INSTI binding site. This data can be found in the supplementary file.

### Protein Secondary Structure assignment

Secondary structural features were identified along the full length of integrase (see Supplementary Materials Table S1). The N-terminal domain (NTD), spanning residues 1-47, mainly consisted of one alpha helix and one 3_10_ helix. These structures may facilitate recognition of viral DNA, which is a function of the NTD.^16^ Meanwhile, the catalytic core domain (CCD) (residues 58-202) had a variety of secondary structures present. The main features were alpha helices and beta strands, but there were some 3_10_ helices, bends and occasional H-bonded turns. Secondary structures found in the C-terminal domain (CTD) (residues 220-270) were by mainly beta strands plus a small number of H-bonded turns. This may be related to the CTD’s multiple accessory functions which relate to other stages in the HIV replication cycle, for example interacting with HIV reverse transcriptase, RNA binding and chromatin remodelling.^17^ It should be noted that certain sections of the integrase protein, namely residues 229-235 and 269-288 were not represented in the crystal structure (PDB: 8fn7^31^).

### Solvent Accessibility

The surface area accessible to solvent of the protein/residues varied from 0 Å^2^ to 210 Å^2^ overall (see Supplementary Materials Table S1) but some parts of the protein structure posed a problem for calculating solvent accessibility. For example, R228 was solved pointing inwards which matched the 0.02Å^2^ solvent accessibility calculated and seemed to form part of a loop region (PDB: 6u8q^32^, a cryoEM structure). However, it was likely to be flexible with a high B-factor of 480.81 as it was solved by cryoEM at a resolution of 4.67Å, making the solvent accessibility calculation for this residue unreliable.

### Mutation frequency

A total of 24568 HIV1 integrase amino acid sequences were extracted from the Los Almos database.^12^ After calculating mutation frequencies of individual residues across all sequences, the overall mutation frequency by residue is shown in Figure 3, while the full table of information is provided in Table S1 in Supplementary Materials.

**Figure 3.**
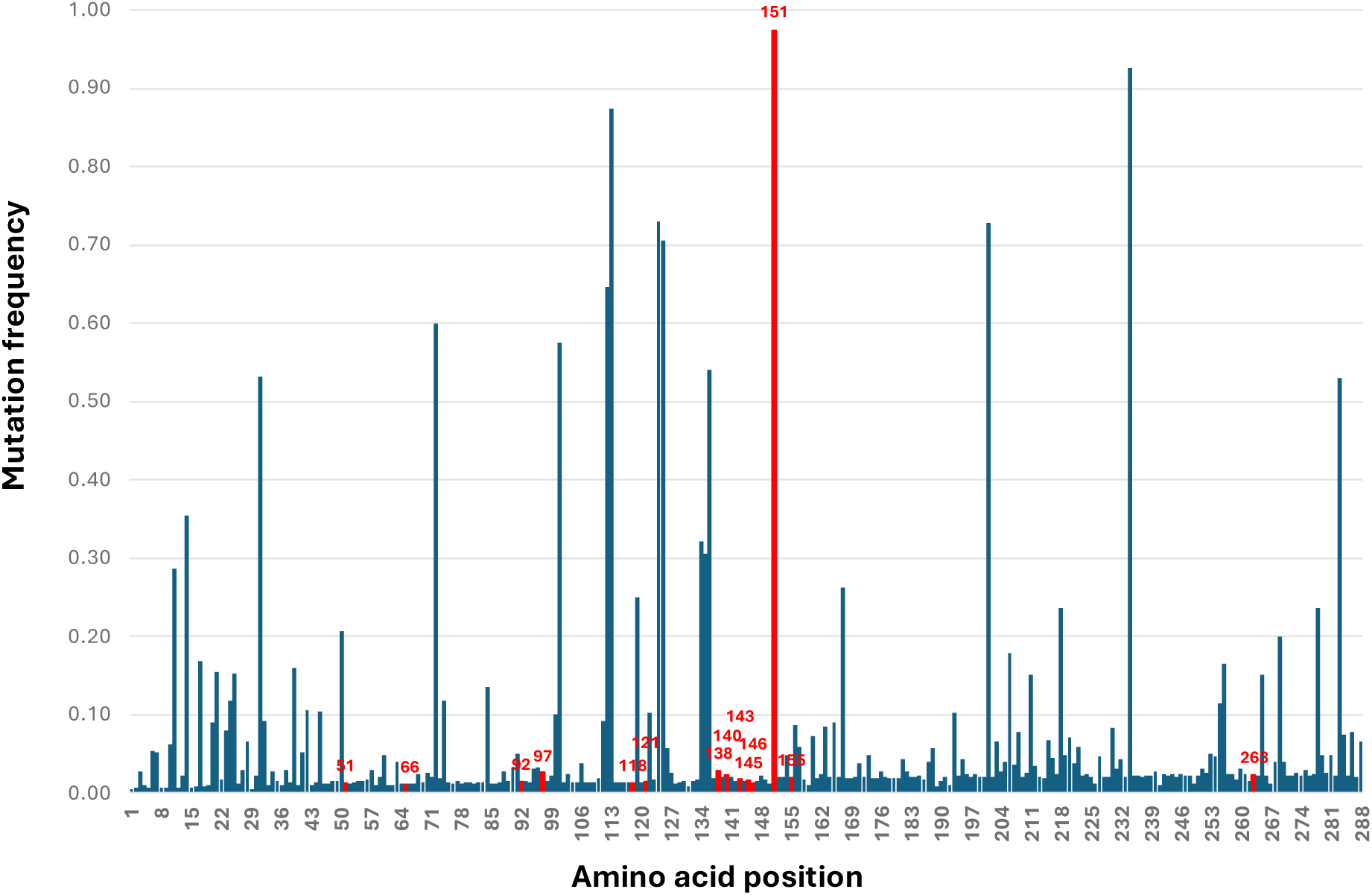
Mutation frequency of all residues in HIV integrase. Amino acid positions highlighted in red corresponds to position of known major RAMs.

From Figure 3, it can be seen that positions corresponding to major RAMs have little or no correlation with mutation frequency. The six positions with the highest mutation frequencies are listed in Table 1.

**Table 1.**
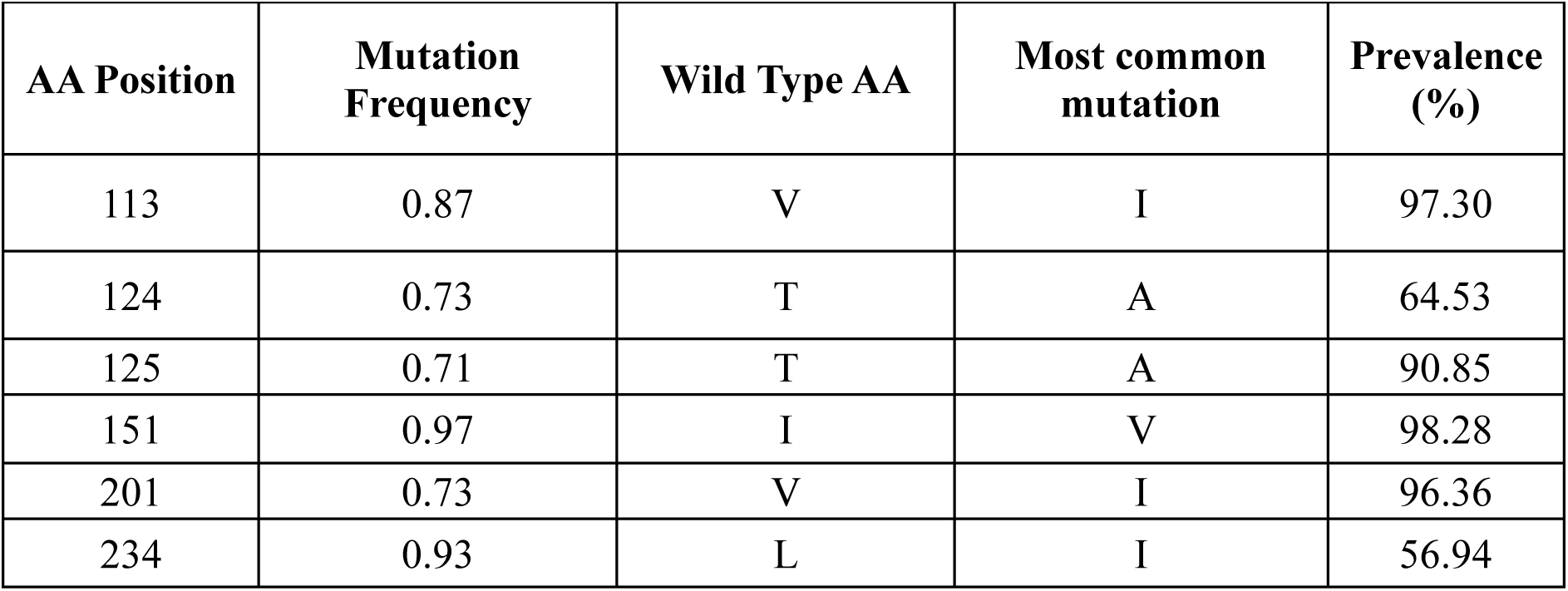
Residues/AA with the highest mutation frequency.

| **AA Position** | **Mutation Frequency** | **Wild Type AA** | **Most common mutation** | **Prevalence (%)** |
| --- | --- | --- | --- | --- |
| 113 | 0.87 | V | I | 97.30 |
| 124 | 0.73 | T | A | 64.53 |
| 125 | 0.71 | T | A | 90.85 |
| 151 | 0.97 | I | V | 98.28 |
| 201 | 0.73 | V | I | 96.36 |
| 234 | 0.93 | L | I | 56.94 |

Prevalence is a measure of the how common a specific mutation is compared to other possible mutations at the same position. It was calculated as follows:

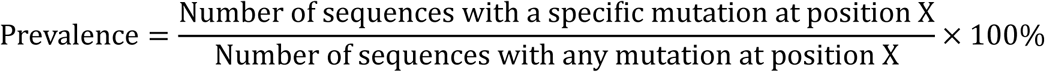

A high mutation frequency (close to 1) means the residue mutates a lot. The prevalence is specific to a particular mutation, and it measures how often it mutates to a particular amino acid if there is a mutation e.g. if V113 mutates to I, 97.3% of the time, giving it a prevalence of 97.36. The high mutation frequencies observed in T124A, T125A, V201I and L234I can be explained by them being common natural polymorphisms found in treatment-naïve patients.^18^ These mutations are not considered RAMs and have little to no effect on INSTI drug resistance.

Residue 151 has an unusually high mutation frequency of 0.97. Interestingly, its most common mutation I151V is not a RAM. The major RAM in this position is I151L^9^ which is very rare, as seen from its 0.04% prevalence. Although residue I151 is part of the drug binding site, I151V has no effect on drug binding or protein function.^19^ I151V is especially common in raltegravir treated patients.^20^ It has been also noted that I151V often appear along with V113I^20^, another mutation with high frequency (Table 2), for unknown reasons. On the other hand, I151L alone can increase the IC_50_ fold change above the resistance threshold for raltegravir and elvitegravir.^21^

**Table 2.** Scores obtained after 10-fold cross validation across a range of *λ* values using the SAGA and LIBLINEAR solvers.

| $\lambda$ | Mean Score ± Standard Deviation | |
| --- | --- | --- |
|  | SAGA | LIBLINEAR |
| 0.01 | $0.94 \pm 0.02$ | $0.93 \pm 0.02$ |
| 0.05 | $0.95 \pm 0.02$ | $0.95 \pm 0.02$ |
| 0.1 | $0.95 \pm 0.02$ | $0.95 \pm 0.02$ |
| 0.5 | $0.95 \pm 0.02$ | $0.95 \pm 0.01$ |
| 1 | $0.94 \pm 0.02$ | $0.94 \pm 0.02$ |
| 5 | $0.90 \pm 0.02$ | $0.90 \pm 0.02$ |
| 10 | $0.87 \pm 0.03$ | $0.87 \pm 0.04$ |

Another interesting discovery from the mutation frequency analysis was that there were many sequences containing identical mutations. Without removing duplicates, over 40000 sequences were extracted from the Los Almos database.^12^ One integrase sequence, for example, was found 347 times (<1% of total sequences) containing the following mutations: K7R, K14R, V54I, V72I, L101I, V113I, I135V, I151V, T218S and V234L. Notably, none of these mutations are RAMs and could be naturally occurring polymorphisms like V113I.

### Model Evaluation

After performing 10-fold cross validation for different values of the strength of regularisation *λ*, and trying different solver algorithms, the scores obtained were as listed in Table 2. Lower *λ* values are slightly favoured over higher ones, suggesting that heavier regularisation may be detrimental to model quality.

From Table 2 neither the *λ* values nor choice of solver have a major impact on prediction accuracy. The choice of solver algorithm appeared to be inconsequential to model performance. The final parameters chosen to proceed with model evaluation were *λ*=0.5, and using the LIBLINEAR solver.

A confusion matrix was produced after testing the model as in Figure 4. It shows that most of our predictions were correct.

**Figure 4:**
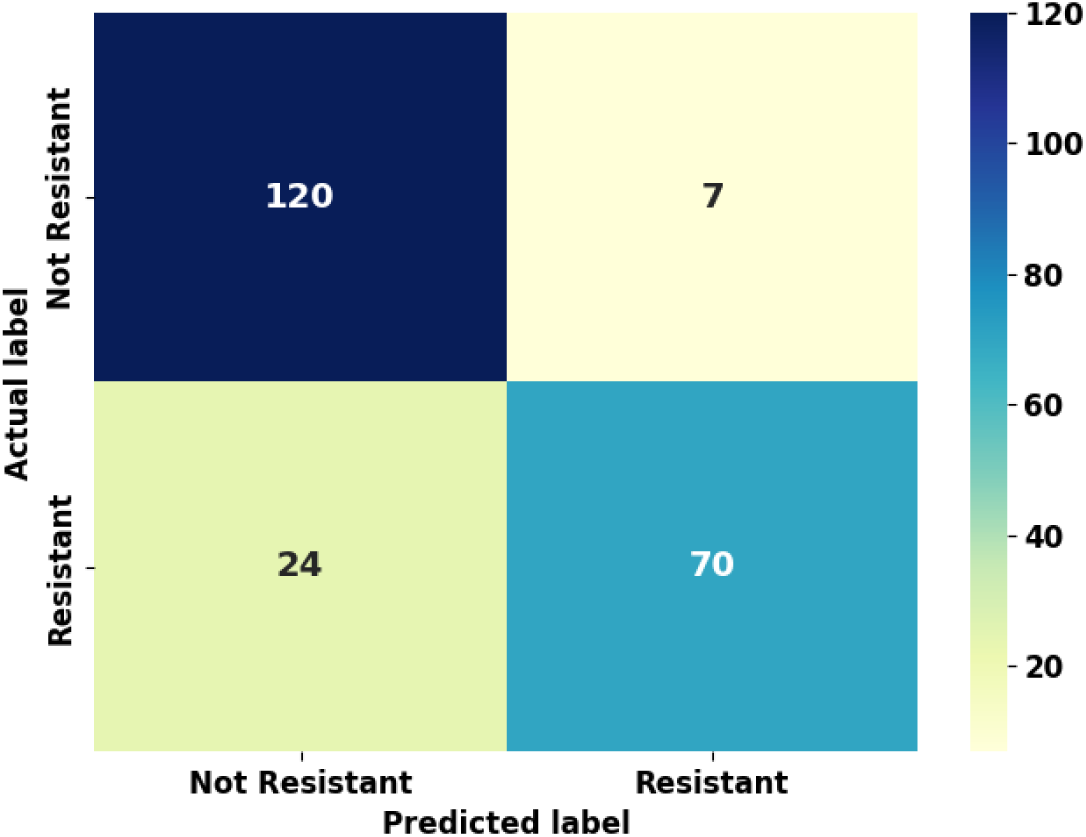
Confusion matrix showing true negative (top left), false negative (top right), false positive (bottom left) and true positive (bottom right). The numbers in the matrix represent the number of sequences in the test set of 221, that fall into each category.

A more detailed classification report was obtained and shown in Table 3.

**Table 3.** Calculated precision, recall and F1 scores for the Classification model.

|  | **Precision** | **Recall** | **F1 score** | **Support** |
| --- | --- | --- | --- | --- |
| **Not Resistant** | 0.83 | 0.94 | 0.89 | 127 |
| **Resistant** | 0.91 | 0.74 | 0.82 | 94 |
| **Macro Average** | 0.87 | 0.84 | 0.85 | 221 |
| **Weighted Average** | 0.87 | 0.86 | 0.86 | 221 |

The “Support” column shows the corresponding number of sequences in the test set of 221 sequences.

Precision, recall and F1-score are different metrics that give insight into model performance.^5^ They were calculated as follows:

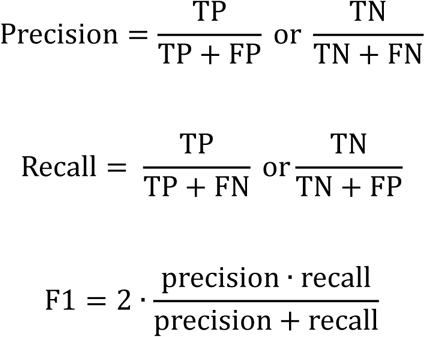

From the model, a ROC curve was also plotted using data from the confusion matrix yielding an AUC of 0.92.

All the metrics above seem to indicate the model is reasonable at predicting drug resistance. Precision was good, with the values for resistant and non-resistant prediction being over 0.8. The recall was slightly low for the resistant group, with a value of 0.74, but was higher in the non-resistant group. For both groups, the F1 score was above 0.8 suggesting a good model quality. The AUC indicated an over 90% chance of correct classification.

### Model Coefficients

After evaluation, the model was fitted, and its coefficients calculated. These measure the weight of each variable and quantify the classification of the prediction. The calculated coefficients, with a non-zero value, are shown in Table 4. A total of 20 residues are predicted that could be highly likely drug resistant. If the absolute value of the coefficient is large, it means the model thinks it is more important in determining resistance. For example, Q148H has the largest absolute value or magnitude of 2.4, therefore it is the most important. The “No” in the RAM column means there are no RAMs associated to that position, however, there might still be mutations.

**Table 4.**
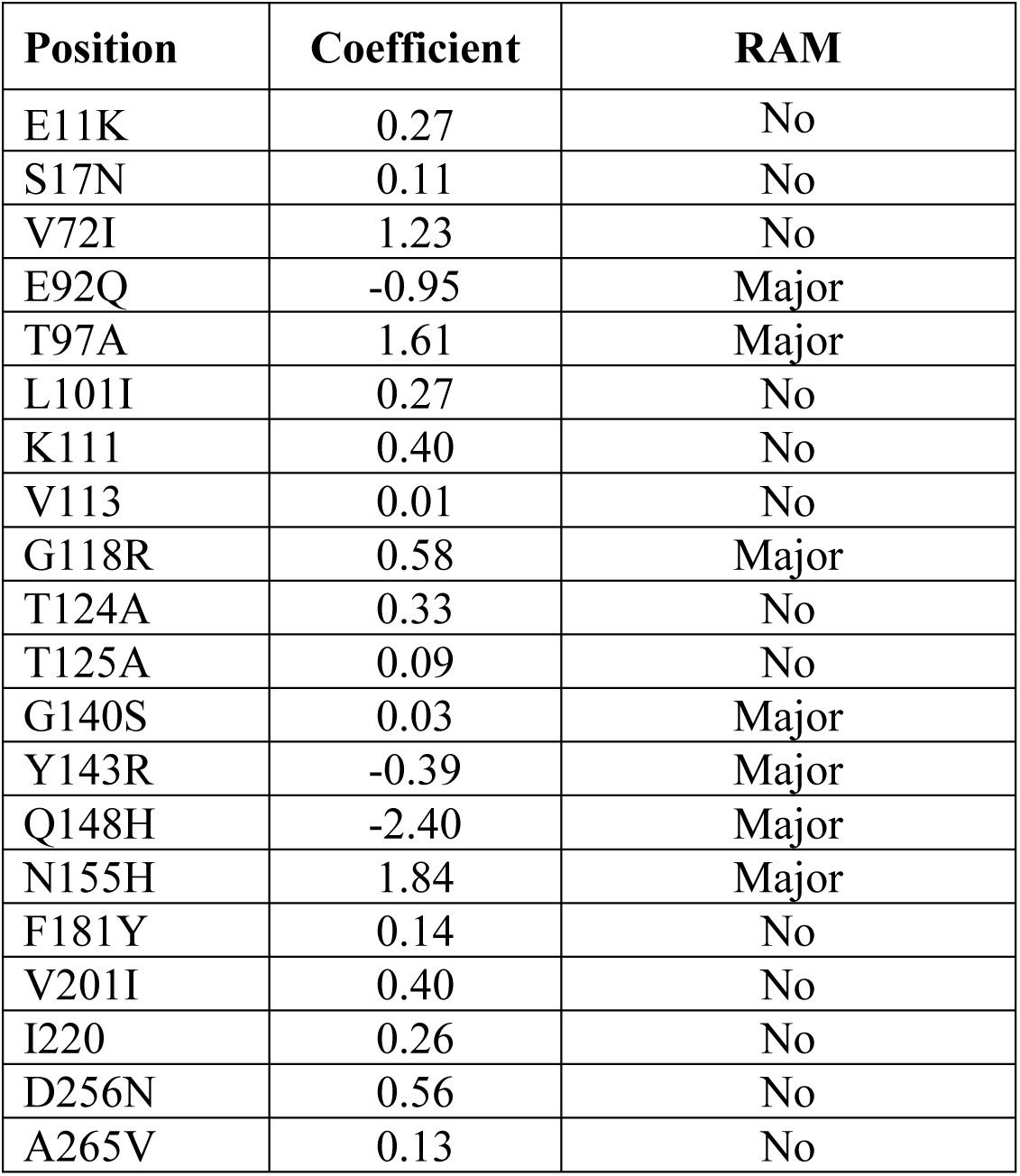
Drug resistance amino acid positions with a non-zero coefficient value.

## DISCUSSION

### Features identified by the model

The model identified 20 mutations regarded as important in the classification of drug resistance as summarised in Table 4. Our model did not find every RAM; although several major RAMs were identified and are labelled accordingly in Table 4. The other identified mutations have some features that seem to be unrelated to drug resistance but could still be important. For instance, while E11K is not considered a RAM, it could prevent formation of a functional tetrameric intasome by changing the negative charge of the glutamic acid^22^ sidechain to positive charge sidechain of the lysine. A similar case could be seen for K186E (not listed in Table 2). Here the positive sidechain of lysine is mutated to a negative charge of glutamic acid. The change of charge can disrupt intasome formation.^35^ However, in our model, it was deemed to be unimportant and given a coefficient of zero. Therefore, the significance of E11K in INSTI drug resistance is uncertain.

S17N, a common polymorphism, has been found to have compensatory effects in correcting abnormalities caused by other mutations,^23^ while V72I is a polymorphism found in patients in the early stages of HIV infection which was tested to have no effect on INSTI resistance^24^.

Although L101I and T124A were not considered RAMs, they were shown to be significantly more frequent in patients treated with raltegravir than in treatment naïve patients.

Furthermore, both mutations have been observed to develop under selection pressure of dolutegravir.^25^ T125A is likely to influence protein-DNA interaction, as T125 was found to form non-bonded interactions with host DNA.^20^ F181Y is a mutation that does not affect viral fitness or drug resistance. The F181 residue forms part of a *π*- *π* interaction along with F185 and W132^26^ (see Figure 5).

**Figure 5.**
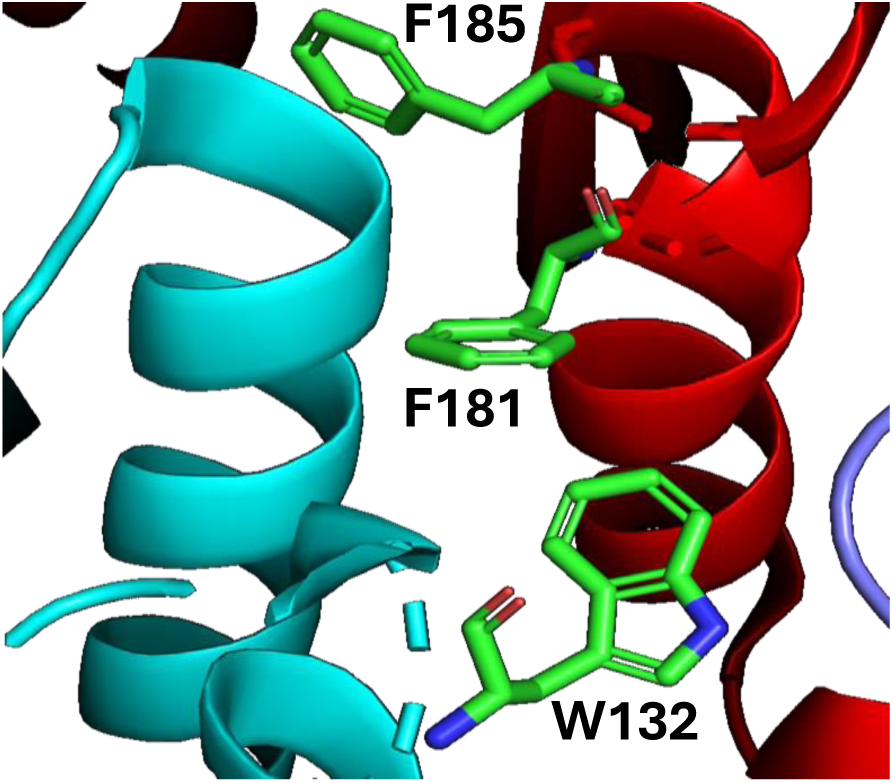
A *π*- *π* interaction is formed by interactions between the residues highlighted in green. Notice that while F181 and F185 are from the same chain shown in red, W132 is from another chain shown in cyan.

This *π*- *π* interaction is crucial for the catalysis of strand transfer. A mutation that removes the benzene ring, meaning removing the capability of forming the *π*- *π* interaction, for instance F181G, results in decreased 3’ processing^26^ while F181Y has no effect as the tyrosine side chain can also form the *π*- *π* interaction.

V201I and A265V were found to be common polymorphism in the database. The V201I substitution has been hypothesised as increasing the binding energy between two chains of integrase.^27^ However, its location in the CTD makes it an unlikely participant in INSTI resistance. D256N is a compensatory mutation that allows binding of integrase to viral RNA.^28^ This mutation provids resistance against inhibitors that block integrase-RNA binding.

In summary, about half of the mutations labelled important are by the machine learning model, major RAMs previously shown to cause INSTI resistance by phenotypic assays. L101and T124A have been found to be mutations induced by INSTI treatment and although they had not been verified as RAMs, they may play a role in resistance. Other mutations labelled as important were either common polymorphisms or had alternative functional importance. Despite their potential importance in other aspects of integrase function, it was difficult to link these other mutations to INSTI resistance. Therefore, the model was somewhat successful in identifying several important mutations but failed to give further insight into how INSTI resistance arose as half of the features deemed important were irrelevant.

### Limitations of the Model

In this investigation, the logistic regression model performed well in cross validation and was therefore likely to succeed in predicting the resistance status of an unseen sequence. While logistic regression has previously been shown to be effective in predicting drug resistance and identifying important features^6^, it has several limitations.

The logistic function assumes a linear combination of features. While convenient to compute, such linear relationships are rarely observed. This may be the main cause of the model’s failure to predict all major RAMs and other important residues as they may not be linearly related. To model more complex relationships, deep learning classifiers such as artificial neural networks should be used. However, as shown in a study using deep learning models in predicting drug resistance^5^, deep learning models trade interpretability for accuracy, resulting in the black box phenomenon mentioned earlier. While such a model would likely be a better fit for the data than generalised linear models, it would be difficult to understand the biological reasonings behind the classifications, making it challenging to justify its results.

Features are considered independently in linear models. However, secondary structure and solvent accessibility both depend on the residue position in the 3D structure. Additionally, residues close to one another are likely to share secondary structure and solvent accessibility.

This type of linkage between features may introduce inaccuracies into the model. Furthermore, minor RAMs had no direct influence on resistance making them difficult to identify using a linear model. This could explain why only major RAMs were identified in the logistic regression model.

A limitation which bottlenecks machine learning in general is the lack of data. The Stanford database only had around 2000 sequences with resistance fold change data available, compared to the over 40000 integrase sequences available on Los Almos database. This is likely caused by the time and cost required for sequencing and performing assays. The shortage of input data heavily impacts model performance and may be reflected in this current investigation.

### Machine learning model comparison with other HIV-1 proteins

Most work done on HIV sequence analysis relating to drug resistance has focused on HIV protease and reverse transcriptase (RT) inhibitors. There are more available drugs in these two classes than for integrases, and more sequence data are available. In a study where deep learning models were used to investigate protease and reverse transcriptase inhibitors, the best performing classifier identified an average of 5 known RAM positions out of the top 20, and the most important feature did correspond to a known RAM position.^5^ This could be contrasted to the current investigation, in which, of all 20 features identified, 7 corresponded to known RAM positions, with two other mutations strongly implicated to be RAMs. From this, a possible conclusion is that a simpler linear model can outperform deep learning models in predicting drug resistance without sacrificing accuracy but with the additional merit of interpretability.

## CONCLUSION

The main difference between the current investigation and previous studies lies in the use of chemical, physical, and biological features such as protein secondary structure, solvent accessibility, and mutation frequency. The inclusion the latter did not improve identification of potential RAMs as the mutation frequencies were rather random and were heavily biased by polymorphisms within certain sample populations. On the other hand, solvent accessibility and secondary structure were helpful in identifying why certain residues may be important, but in many cases it was difficult to relate their functional significance to INSTI resistance.

The common feature relating all the known RAMs identified by the model was their presence in the drug binding site, as 6 out of 7 of the identified RAMs were located in the binding site. This is reasonable given the competitive enzymatic mechanism of INSTIs.

There are other linear models available which have been successfully applied to drug resistance classification problems, with a popular option being support vector machines^2^. Accounting for the strengths and limitations of different types of classification models, the ideal method would be to train the data using different models and compare their performances. Many studies have used different machine learning methods to train their models and resulted in more potential solutions to the problem. We have also attempted various training methods. However, in this investigation, none of the other methods returned meaningful results.

Regarding the assumption of feature independence by linear models, a possible workaround could be to use large language models instead. A large protein language model has been developed to predict antibiotic resistance-associated genes and resistance classification to a high degree of accuracy.^29^ Large language models are much more complex than linear models and specifically fall within the category of transformer models, which emphasise on context and how different features relate to one another. In the same way that words at various positions in a sentence relate to one another, so potentially related residues can be found in different position in a protein sequence. Identifying these linkages could be the key next step in improving prediction accuracy and identification of new RAMs.

For a compromise between model complexity and consideration of potential links between residues, it is perhaps possible to group residues on the protein 3D structure into local clusters, since proximal residues are more likely to interact with one another. To identify these clusters, a probe (for instance a 10Å x 10Å x 10Å cube) could be moved around in 3D space and residues that fall within the probe’s range be considered a cluster. This method relies on the assumption that only proximal residues are important, and therefore overlooks the distal interactions. Excluding distal residues of course significantly reduces model complexity, and their inclusion would not necessarily improve the results.

## DATA AVAILABILITY

The method was implemented using Python 3.7. The Python Integrated Development Environment used was PyCharm. The code for the method was made publicly available on GitHub: https://www.github.com/ethanchuicy/insti_resistance_model

## SUPPLEMENTARY MATERIALS

The file supplementary.csv contains the mutation frequency, solvent accessible surface area, secondary structure, and residues defining the binding site of each residue in integrase, is available for download.

